# Weakly supervised contrastive learning predicts tumor infiltrating macrophages and immunotherapy benefit in breast cancer from unannotated pathology images

**DOI:** 10.1101/2023.04.30.538851

**Authors:** Guobang Yu, Yi Zuo, Bin Wang, Hui Liu

**Affiliations:** College of Computer and Information Engineering, Nanjing Tech University, 211816, Nanjing, China; Department of Cardiothoracic Surgery, the Third Affiliated Hospital of Soochow University, Changzhou, 213110, Jiangsu, China

**Keywords:** Whole Slide Image, Contrastive Learning, Tumor Immune Microenvironment, Macrophages, Biomarker Gene, Computational histopathology, Immunotherapy

## Abstract

The efficacy of immune checkpoint inhibitors is significantly influenced by the tumor immune microenvironment (TIME). RNA sequencing of tumor biopsies or surgical specimens can offer valuable insights into TIME, but its high cost and long turnaround time seriously restrict its utility in routine clinical examinations. Several recent studies have suggested that ultra-high resolution pathology images can infer cellular and molecular characteristics. However, studies on revealing TIME from pathology images are still limited.In this paper, we proposed a novel weakly supervised contrastive learning model to deduce tumor immune microenvironment features from whole slide images (WSIs) of H&E stained pathological sections. The high-resolution WSIs are split into tiles, and then contrastive learning is applied to extract features of each tile. After aggregating the features at the tile level, we employ weak supervisory signals to fine-tune the encoder for various downstream tasks. Comprehensive downstream experiments on two independent breast cancer cohorts and spatial transcriptomics data demonstrate that our computational pathological features accurately predict the proportion of tumor infiltrating immune cells, particularly the infiltration level of macrophages, as well as the immune subtypes and biomarker gene expression levels. These findings demonstrate that our model effectively captures pathological features beyond human vision, establishing a mapping relationship between cellular compositions and histological morphology, thus expanding the clinical applications of digital pathology images.

## I. Introduction

**B**REAST cancer is a type of malignant tumor that poses a significant global threat to women’s lives, its incidence and mortality rates show an overall upward trend. In 2020, breast cancer accounted for approximately 12.5% of newly diagnosed cases and 6.92% of total cancer-related deaths [1]. Despite considerable advancements in the treatment of breast cancer in recent years, remains a significant threat to women’s lives globally. Although immune checkpoint inhibitors (ICIs) [2] have shown promising efficacy in various solid tumors [3], their benefits are limited to a small subset of breast cancer patients.

Numerous studies has confirmed that the efficacy of ICIs in solid tumors is influenced by the tumor immune microenvironment (TIME) [4]. TIME refers to the degree and distribution of immune cell infiltration within the tumor,and plays a crucial role in tumor proliferation, immune evasion mechanisms, and the effectiveness of immunotherapy. To assess the composition of immune cells in the tumor microenvironment, quantitative analysis using RNA sequencing (RNA-seq) data [5] are typically conducted using tools such as CiberSort [6], ssGSEA [7] and ESTIMATE [8]. “However, the high cost and long turnaround time of RNA-seq not only impose a financial burden on patients but also potentially lead to missing the optimal treatment window. There is an urgent need for low-cost and rapid methods to complement TIME quantification.

In recent years, deep learning [9] has achieved tremendous success in the field of computer vision. For example, for the image classification task on ImageNet [10]–[12] dataset the deep convolutional neural network (CNN) model has achieved better performance than human [13]. As a result, some researchers have applied deep learning with high-resolution histopathological images for automatic tumor diagnosis and subtyping. Wang et al. developed an improved HD-Staining model based on Mask-RCNN to automatically segment and identify cell nuclei in lung adenocarcinoma [14]. Campanella et al. proposed a weakly supervised learning model that performed well in tumor diagnosis based on WSIs [15]. Thomas et al. developed a multi-instance learning model [16] that can distinguish between benign tissue and metastatic breast cancer [17]. Chen et al. proposed a method based on deep learning that use annotation-free pathology image to distinguish lung cancer subtypes [18]. Ming et al. proposed the CLAM model that aggregates tile features using an attention mechanism to accurately classify subtypes of renal cell carcinoma and nonsmall cell lung cancer [19]. Moreover, a few studies went further step to infer molecule-level features, such as genetic alteration and gene expression. For instance, Kather et al. reported a deep learning-based model that predicts microsatellite instability in gastrointestinal cancer from WSIs, and showed that pathological image features can reveal gene mutations, gene expression characteristics, and standard biomarkers, elucidating and quantifying the genotype-phenotype relationship in cancer [20], [21]. Hu et al. constructed SpaGCN, which combines gene expression, spatial location, and pathology image to identified spatial domains and spatially variable genes [22]. Schmauch et al. developed the HE2RNA deep learning model to predict tumor RNA-seq expression from WSIs [23].Several studies utilize pathological images to investigate the tumor immune microenvironment. Seung-Yeon et al. leveraged pathology images to characterize the tumor immune microenvironment in colorectal cancer [24]. Saltz et al. used a deep learning model to demonstrate the infiltration state of tumor-infiltrating lymphocytes (TILs) and generated spatial maps to establish prognostic risk score [25]. These studies have demonstrated the promising potential of pathology images in disentangling TIME. However, these methods rely on large-scale of manual annotations of various types of immune cells by experienced pathologists, which are often not available in practice.

Self-supervised learning [26] eliminates the necessity for human-annotated labels by generating pseudo-labels autonomously from the data. This approach uses unlabeled data to learn robust features or representations for various downstream tasks. Our team has utilized contrastive learning to extract expressive features from pathology images and predict differentially expressed cancer driver genes [28]. In this paper, we integrated contrastive learning and weakly supervised learning [29] to investigate the tumor immune microenvironment in breast cancer. We initially partitioned high-resolution whole-slide images (WSIs) into tiles and excluded tiles with white backgrounds or insufficient tissue. Next, we used contrastive learning to extract tile features and then utilized attentive pooling [30], [31] to aggregate tile-level features to construct the slide feature. We used weak supervisory signals to fine-tune the slide-level features and applied it to various downstream tasks, including inferring proportion of tumor immune infiltration cells, immune subtyping, expression level of biomarkers and patient survival. Experiments demonstrate that our model effectively predicted the tumor infiltration levels of 22 types of immune cells well. Notably, it attained superior performance for predicting three types of macrophages and the biomarker gene expression levels. For the discrimination of six immune subtypes, our method also achieved a 0.8 ROC-AUC value. Moreover, the attention scores of tiles generate heatmaps that are indicative of tumor regions and spatial localization of biomarkers. For instance, the high-attention areas of the heatmap generated from the tumor diagnosis task were highly consistent with tumor regions annotated by an experienced pathologist with 10 years of experience. The spatial expression pattern of biomarker genes by spatial transcriptomics also validated the capacity of our method in spatial localization of gene expression. These findings suggests that our method proficiently established the genotype-phenotype associations, significantly promoting the potential of digital pathology in clinical applications.

## II. Materials and methods

### A. Whole slide image and annotations

All hematoxylin and eosin-stained fresh-frozen tissue images of the TCGA-BRCA cohort were downloaded from the Genomic Data Commons data portal. All whole slide images were scanned st 20x magnification. This cohort comprises 1,821 WSI collected from 1,094 breast cancer patients, including 1,577 samples from cancer patients and 244 samples from normal patients. The slide-level diagnosis (tumor or nontumor) provided by THE TCGA database were used as ground truth for classification labels. The slides were downloaded from http://portal.gdc.cancer.gov.

An external validation cohort CPTAC-BRCA data were obtained from the TCIA database, which included 344 WSI (20x magnification) from 121 patients. The slides were downloaded from http://faspex.cancerimagingarchive.net.

A pathologist with more than ten-year experience was asked to manually annotated a few slides. The annotated slides were used for the validation of spatial localization of tumor and necrosis regions.

### B. Spatial transcriptomic data

We obtained spatial transcriptomic data of a breast cancer specimen using the Visium Spatial Gene Expression protocol from 10x Genomics. The fresh frozen samples of invasive ductal carcinoma were obtained from BioIVT. This particular sample consisted of 4,898 spots with an average of 3,654 genes and 47,223 reads per spot. The ST-seq data and pathology image can be accessed at http://www.10xgenomics.com. To construct the spatial landscape of gene expression, we utilized the 10x Loupe Browser, a desktop application that provides interactive visualization capabilities for various 10x Genomics solutions. This allowed us to visualize the spatial landscape of specific biomarker gene expression and overlay it onto the pathology image.

### C. WSI preprocessing

Since pathological images are too high-resolution so that they cannot be directly fed into a deep neural network. To overcome this challenge, the WSIs are divided into smaller tiles. First, we utilized the Otsu algorithm [32] to select regions containing enough tissue cells.Specifically, each slide is loaded into memory using OpenSlide and transformed from RGB to the HSV color space. A binary mask of the tissue area (foreground) is created by applying threshold of 8 to the saturation channel in the HSV color space, ensuring smoothing edges and performing morphological closure to filling small gaps and holes. This helps separates the WSI into tissue and non-tissue regions. After segmentation, each WSI is split into 256x256 pixel tiles within the foreground region and white image tiles and white image tiles with insufficient tissue (pixel background value*>*200) are discarded. To enhance access efficiency, only the coordinates and metadata of each tile are stored in hdf5 format.

### D. TIME estimation based on RNA-seq data

To estimate the abundance of tumor infiltrating lymphocytes, we downloaded RNA-seq data of the samples with matched pathology images from theGenomic Data Commons data portal. The next step involved using CIBERSORT [6], a deconvolution method that characterizes the cell composition of complex tissue, to compute the abundance of 22 immune cell types from the RNA-seq data.

### E. Contrastive learning-based computational pathology framework

Our learning framework comprises three stages: image preprocessing, contrastive learning-based pre-training, and downstream tasks. The schematic diagram is shown in Figure 1. Firstly, high-resolution pathology images were partitioned into 256*256 px tiles, with white backgrounds and tiles without enough tissue content removed. In the second stage, contrastive learning (MOCO v2, AdCo [33]) was used to extract features from each tile, the backbone encoder used for feature extraction is ResNet-50. In the third stage, the features extracted from the tiles are aggregated using attentive pooling and the model was fine-tuned using weak supervisory signals from downstream tasks.

**Fig. 1:**
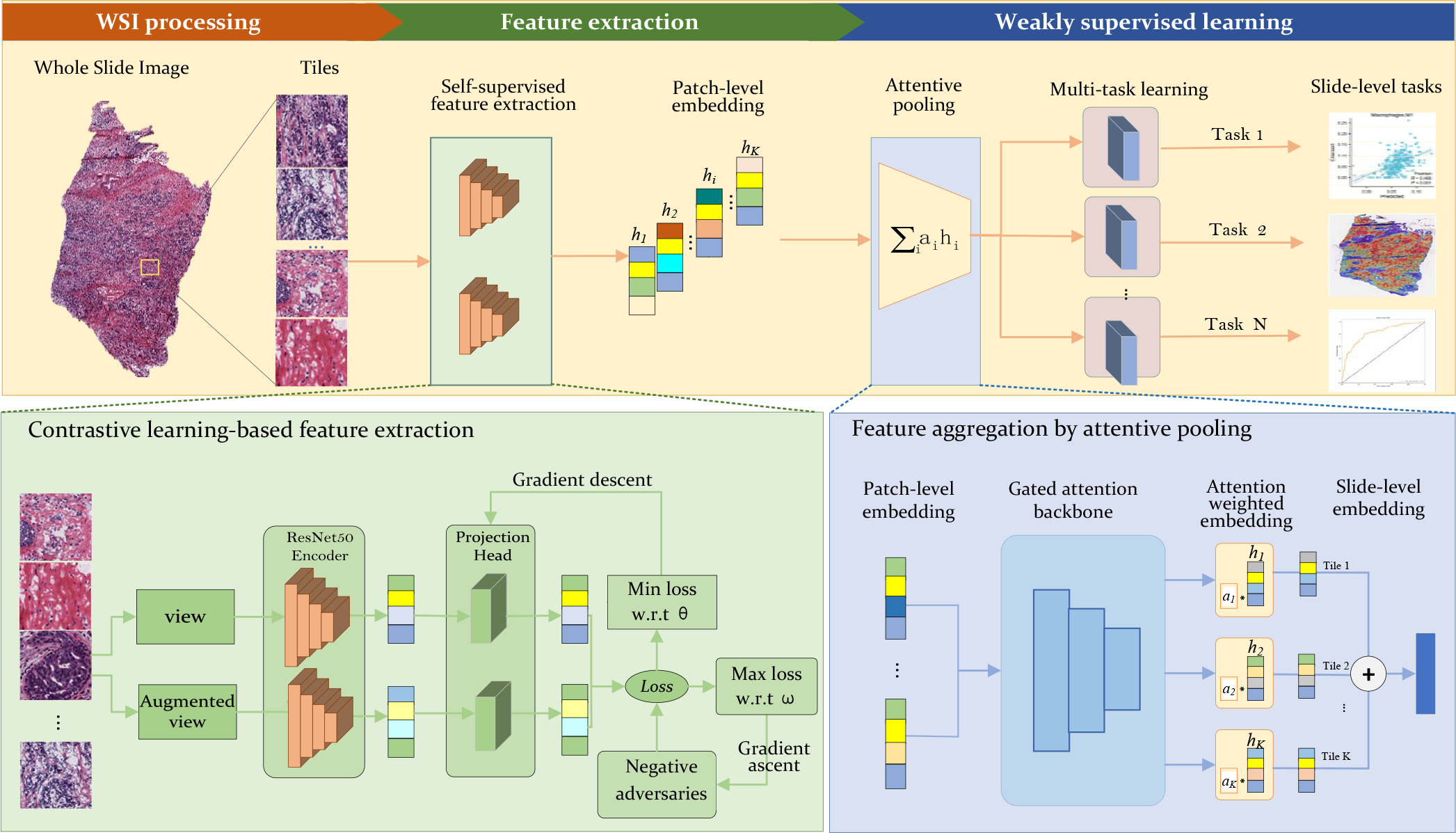
Our learning framework consists of three stages: image preprocessing, pre-training through contrastive learning, and weakly supervised learning for downstream tasks. Initially, high-resolution pathology images were split into 256x256-pixel tiles, with tiles containing white backgrounds or insufficient tissue content being removed. In the second stage, contrastive learning was used to extract features from each tile, with ResNet-50 serving as the backbone encoder. Finally, tile features were aggregated through attentive pooling and adapted to downstream tasks with weak supervisory signals.

For feature extraction, we tested two types of selfsupervised learning, adversarial contrastive learning (AdCo) [33] and masked autoencoder (MAE) [34], and compared their performance. Contrastive learning brings the original image closer to its positive partner (e.g. augmented image) while keeping it as far away from negative samples in each batch as possible, thereby forcing the encoder to capture the essential features of each tile and obtain expressive latent representation. The masked autoencoder was trained to reconstruct missing pixels in an input image by masking random patches of the image. Our previous work [26] and the experiments conducted in this study (see Section 4) have confirmed that contrastive learning-based pre-training significantly improved the predictive performance in downstream tasks.

Unlike previous studies that used single-task learning, such aseach gene or mutation had its own dedicated model, we employed multi-task learning in downstream tasks. Multiple tasks share the representation of a shallow network. The related tasks communicate with each other during training, helping in forcing the encoder to learn more generalizable features, thereby enabling our model to obtain stronger inductive bias and robustness.

### F. Contrastive learning-based tile feature extraction

We employed self-supervised contrastive learning on a large collection of unlabeled tiles to derive their latent representations. We utilized the Adversarial Contrastive learning (AdCo) [33] algorithm for pretraining. AdCo operates by constructing a set of negative examples that are challenging to distinguish from positive queries.

The positive pairs were generated through the use of data augmentation techniques such as image rotation, random cropping, and color transformation. Denote the input tile by *x*_*i*_, which is transformed into a latent representation *h*_*i*_ =f(*x*_*i*_) through an CNN encoder *f*. This representation is then projected onto an embedding *q*_*i*_ via multiple fully-connected layers. A queue is initialized to cache the latent representations of the tiles as negative samples. The network parameters *θ* are trained through contrastive learning to distinguish between the query sample *q*_*i*_ and its positive counterpart from a set of negative samples *K*. The contrastive loss is defined as:

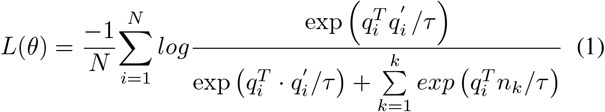

where *q*_*i*_ and 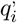 represent the embeddings of a parir of positive samples, *n*_*k*_ refers to the embedding of a negative sample, and *τ* is the temperature parameter. AdCo [33] is inspired by adversarial learning and aims to generate challenging negative samples that are difficult to distinguish from query samples. It extends the memory bank used to store negative sample embeddings by treating negative samples as learnable weights. This allows the learned negative samples to closely track changes of the learned embedding representations.

In our implementation, we employed ResNet50 as the backbone network and its weights were initialized using those learned on ImageNet. During the training process, we adopted SGD as the optimizer. The learning rate of the backbone network is set to 0.03. Weight decay is set to 0.0001 and momentum is set to 0.9.

### G. Gated-attention pooling of tile-level features

Since the downstream task relies on slide-level features, we need to aggregate tile-level features. Rather than max pooling and average pooling, we employ gated attention pooling to aggregate the tile features. Suppose a slide is divided into *N* tiles we compute the slide-level feature as below:

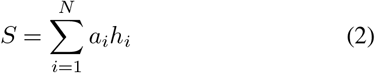

where *a*_*i*_ represents the learnable parameter that indicates the importance of the *i*-th tile to a specific downstream task. Formally, *a*_*i*_ is defined as follows:

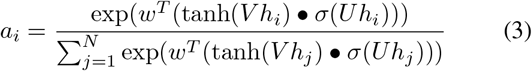

where *U* and *V* were both learnable parameters, represented an element-wise multiplication, *σ* represented the sigmoid activation function, and *w*^*T*^ is used to transfer the feature vector into a raw attention score. The key difference between gated-attention pooling and max/mean pooling is that it introduces learnable parameters that are updated iteratively during training, making the model highly flexible and adaptable to different downstream tasks. More importantly, the learned attention weights reflect the importance of tile-level features to the downstream task, which makes our model interpretable.

### H. Weakly supervised learning for downstream tasks

We applied weakly supervised learning to leverage slide-level features for various downstream tasks, including both classification and regression tasks.

The predictions of tumor infiltrating lymphocytes and biomarker gene expression levels were formulated as regression tasks. Taking as input the extracted slide features, we used a multi-task learning model with a fully-connected layer and an output layer to predict the actual values. For instance, the output layer had 22 nodes corresponding to proportions of 22 types of tumor infiltrating lymphocytes. In our practice, we have actually tested multiple hidden layers, and found that a single hidden layer could achieve superior performance in the regression task. Denote by *M* the number of patients, *y* and *y*^*′*^ the true and predicted values, we used the mean squared error as the loss function:

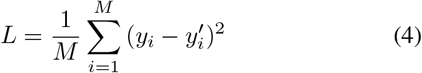

We formulated the tumor diagnosis, tumor immune subtyping and potential immunotherpay benefit as classification tasks. We employed a fully-connected layer plus a softmax layer as the prediction model that took the slide feature as input. The cross-entropy was used as the loss function for the classification task. Denote by *y* and *y*^*′*^ the true and predicted labels, the loss function is as below:

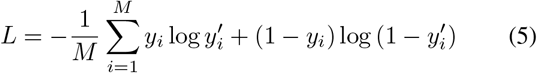

During the training process, we used the Adam [35] optimizer with a weight decay of 0.0001. A decay strategy was applied to dynamically adjust the learning rate. The learning rate was multiplied by 0.1 every time the model trained for 10 epochs.

## III. Results

### A. Weakly supervised learning accurately predict TIME

We first tested the performance of the model in predicting TIME on the TCGA-BRCA cohort. The TCGA-BRCA cohort contains 1,821 WSIs, including 1,577 tumor and 244 normal samples. We formulated this problem as a regression task. The infiltration levels of 22 types of immune cells were determined using the CiberSort tool applied to matched RNAseq data and used as the ground truth. A multilayer perceptron (MLP) model with single hidden layer was constructed, which accepted the 1024-dimension slide-level features as input and had 22 output nodes, each corresponding to specific immune cell type. The Pearson correlation coefficients was used as performance evaluation metrics.

Figure 2 (a) displays the correlation coefficients achieved by our model for 22 immune cell types. We found that the majority of immune cells could be effectively predicted. The average coefficient of 22 types immune cells was 0.30, with 90 percent achieved statistically significant results (pvalues *<*0.01). More interestingly, we observed that higher degree of immune cell infiltration, better accuracy achieved by our model, as shown in Figure 2 (b). It is postulated that an increase in the infiltration of a particular immune cell type results in more pronounced histological morphology in pathological images, thereby facilitating our model to capture features associated with that specific immune cell type. By carefully checking the poor predicted immune cells revealed that the presence of a substantial number of zero values in actual values was the underlying cause. For example, dendritic cell resting, gamma T cells and neutrophil cells contained 76%, 86%, and 61% zero values, respectively. Too many zero included in the actual values led to a collapse in inductive bias. Upon excluded these immune cells with a high proportion of zero values, the average correlation coefficient increased to 0.35. The results strongly supported that our method captures features reflecting the cellular morphology from pathological images. In addition, we compared the performance achieved by using the WSIs with 20x and 40x magnification, and obtained comparable performance (Figure S2(a)). We thus used the 20x magnification slides in subsequent experiments.

**Fig. 2:**
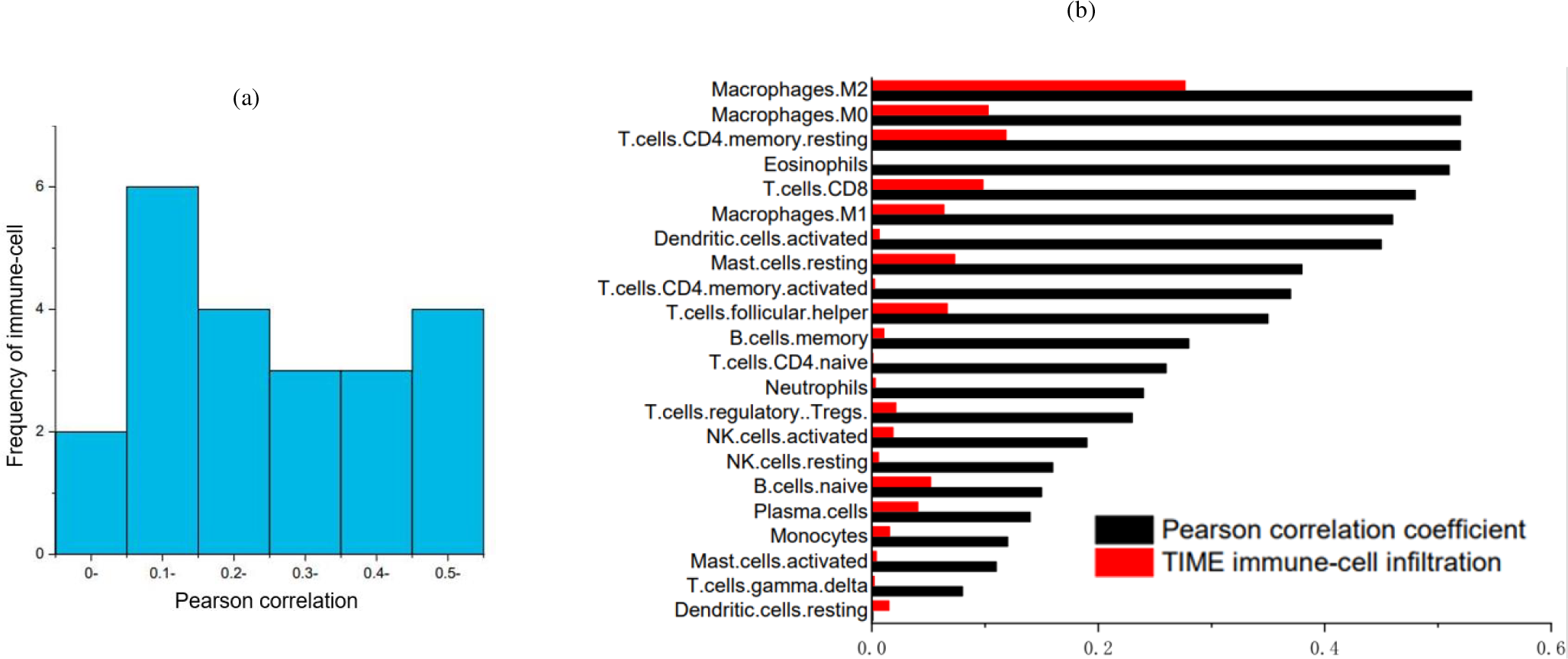
Computational pathological feature accurately predict tumor immune microenvironment. (a) Histogram of the Pearson correlation coefficients verified that the infiltration level of almost all types of immune cells are significantly predicted. (b) The predictive performance measured by Pearson coefficients showed overall correlation to the actual infiltration level of immune cells.

**Fig. 3:**
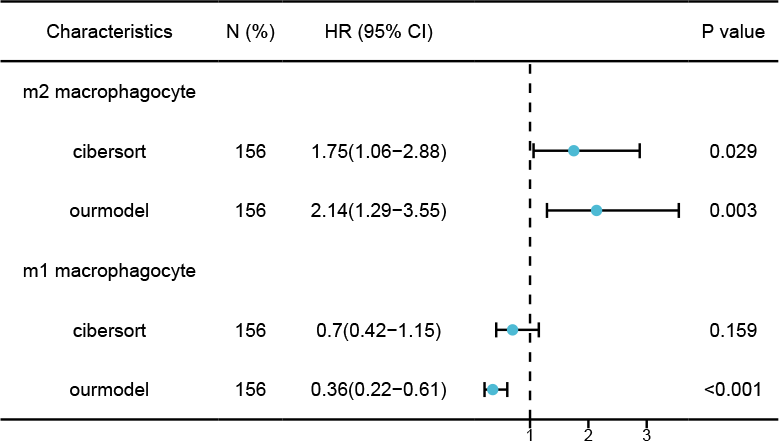
Forest plot reflected the prognostic value of the M1 and M2 macrophage subtypes, and our predicted infiltration level based on pathology images exhibited comparable and even better statistical significance than their actual values.

### B. Accurate prediction of macrophage infiltration level

To further validate the effectiveness of our model, we further explored its predictive performance on macrophages. Macrophages are a ubiquitous immune cell subpopulation that may accounts for over 50% of immune cells in the tumor microenvironment [36]. In breast cancer, macrophages differentiate into M1 and M2 subtypes [37]. In general, M1 macrophages are tumor-resistant due to intrinsic phagocytosis and enhanced antitumor inflammatory reactions. In contrast, M2 macrophages are endowed with a repertoire of tumor-promoting capabilities involving immuno-suppression, neovascularization, as well as stromal activation and remodeling. Our model demonstrated correlation coefficients of 0.466 (p*<*0.001) and 0.502 (p*<*0.001) for M1 and M2 macrophages, respectively. In addition, we employed Cox univariate regression to investigate the influence of these two macrophage subtypes on patient prognosis and compared the predicted values with the actual values obtained from RNA-seq. As depicted in Figure III-A, the forest plot reflected the opposite functions of the two macrophage subtypes. Remarkably, the predicted values exhibited greater statistical significance than their actual counterparts for both M1 cells (p*<*0.001 vs p*<*0.159) and M2 cells (p=0.003 vs p*<*0.029).

### C. Effective prediction of biomarker gene expression levels

To test whether our model could capture more fine-grained information than cells, we employed computational pathology features to predict the expression levels of 11 biomarker genes such as CD40, CD80, and PGK1. These genes serve as common markers of immune cells and are closely associated with prognosis of breast cancer patients [32], [38]. By utiling gene expression levels derived from RNA-seq data as ground-truth values and formulating it as a regression task, we constructed a MLP model with a single hidden layer to predict gene expression using slide-level features as input. We employed multi-task learning to simultaneously predict the expression levels of multiple genes. Our experimental results demonstrated that the predicted marker gene expression levels were highly consistent with the experimental values, with an average correlation coefficient of 0.4 (Figure 4 (a)). For instance, CD80 and CD86 are significant indicators of clinical pathological grading and prognosis and are promising targets for immunotherapy [35]. Our model achieved coefficients of 0.52 and 0.59 for these two marker genes, respectively. Additionally, CD74 (r=0.53) is associated with class II major histocompatibility complex and plays a crucial role in regulating antigen presentation during immune responses [39]. PGK1 (r=0.49) regulates the expression of pro-inflammatory cytokines such as IL-1 and IL-6 in macrophages [40]. These findings suggested that our model has successfully learned fine-grained features and established the connection between genotype and phenotypic feature.

**Fig. 4:**
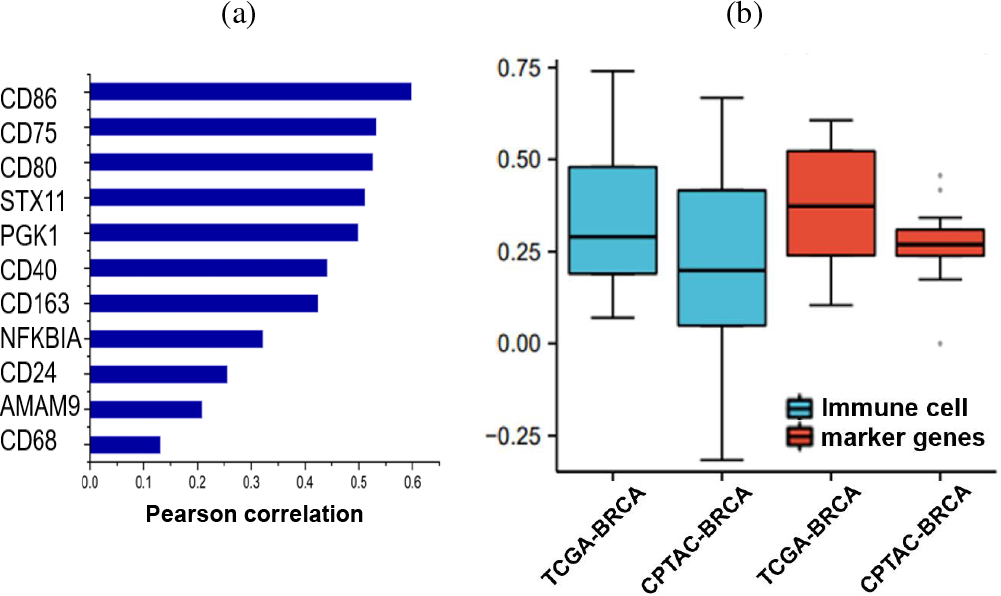
Effective prediction of the tumor immune infiltration levels and expression level of cell-type biomarker genes by pathological feature. (a) Histogram displays the Pearson correlations between the predicted and true values. (b) Statistical boxplots of the Pearson correlations between the actual and predicted immune cell propositions and macrophage marker gene expression levels on TCGA-BRCA and CPTAC-BRCA cohorts. Of note, our model is trained on the TCGA-BRCA cohort and then used for prediction on CPTAC-BRCA cohort.

### D. External independent cohort validated model generalizability

To further validate the robustness of our proposed model, we used an independent cohort from CPTAC-BRCA [41] project to evaluate the model performance in predicting immune cell infiltration and biomarker gene expression levels. It is particularly noteworthy that we did not train and test within the independent dataset, but rather tested on this cohort after model training on the TCGA-BRCA dataset. This strongly validated that our model can capture real features from a large number of pathological images rather than simply remembering the training samples. The CPTAC-BRCA cohort included 344 WSIs of 121 patients, the immune cell infiltration level and biomarker gene expression levels were derived from matched RNA-seq data. As shown in Figure 4 (b), our model achieved an average correlation coefficient of 0.23 for 22 types of immune cell. For the 11 biomarker gene expression levels, our model achieved average correlation coefficient 0.28(see Figure S1(A,B)). The experimental results on the independent cohort strongly validated that our proposed model has solid generalizability.

### E. Computational pathological features inferred tumor immune subtypes and potential immunotherapy benefit

Tumors can be stratified into three fundamental immune subtypes based on the degree of immune cell infiltration within the tumor microenvironment: inflamed, excluded, and desert [42]. The immune-inflamed tumor, also known as “hot tumors”, are characterized by elevated infiltration of immune cytotoxic cells, activation of the interferon-*γ* signaling pathway, and an abundance of M0 and M1 macrophages. Both the immune-excluded and immune-desert tumors may be classified as “cold tumors”. The excluded subtype is distinguished by the presence of CD8+ T lymphocytes at the invasive margin that are incapable of effectively infiltrating the tumor. In contrast, the desert subtype is characterized by the paucity of CD8+ T lymphocytes within or surrounding the tumor. The immune checkpoint inhibitors demonstrate an augmented response rate in hot tumors, whereas the response rate in desert tumors is markedly diminished.

Based on the immune cell infiltration levels derived from RNA-seq data, we utilized the k-Means algorithm to cluster 1,094 cases from TCGA-BRCA cohort into three distinct clusters and assigned them to the excluded, inflamed, and desert immune subtypes. As shown in Figure 5 (a,b), clusters 0 and 2 exhibited high levels of M2 macrophages, resting NK cells, and neutrophils. These cells are generally considered to facilitating tumor cell proliferation, extravasation, and migration and are associated with unfavorable prognosis. In contrast, cluster 1 tumors were infiltrated by relatively high levels of T cells and M1 macrophages that produce proinflammatory cytokines and reactive oxygen intermediates to kill tumor cells.

**Fig. 5:**
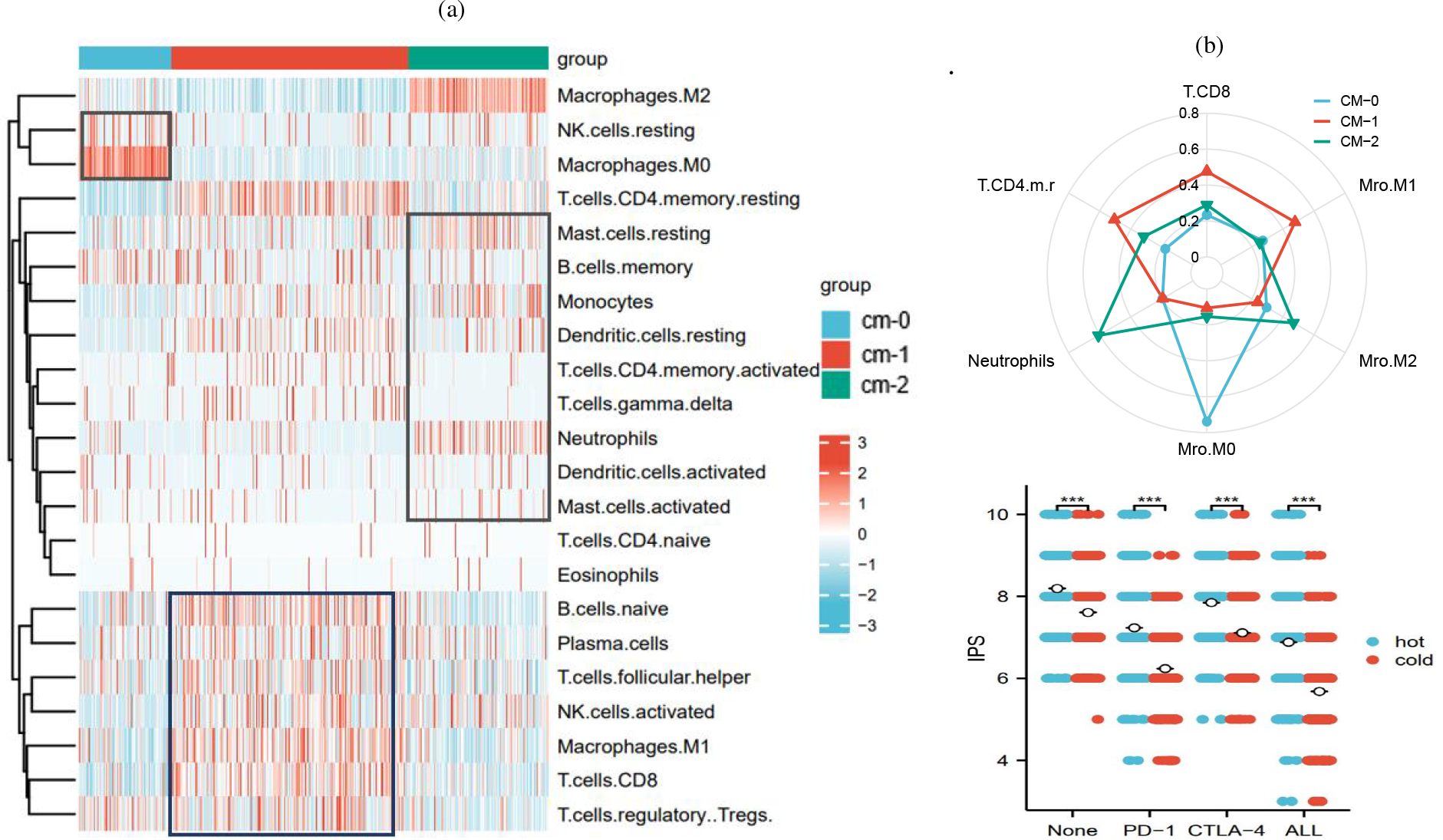
Histopathological features reveal tumor immune subtypes and potential immunotherapy benefit. (a) Heatmap of three clusters generated by the k-means clustering on 21 types of tumor immune infiltrating cells. (b) Radar plot of the tumor immune infiltration level of CM0-2 (uppper) and the IPS scores between the hot and cold tumour groups.

After assigning the immune subtype to each case, we used pathological image features to make predictions. The results are shown in Figure 6 (a), our model obtained an average ROC-AUC value of 0.76. If the immune subtype is simplified into two categories of “cold” and “hot” tumors [43], the predictive model based on pathological images attained 0.75 accuracy and 0.82 ROC-AUC.

**Fig. 6:**
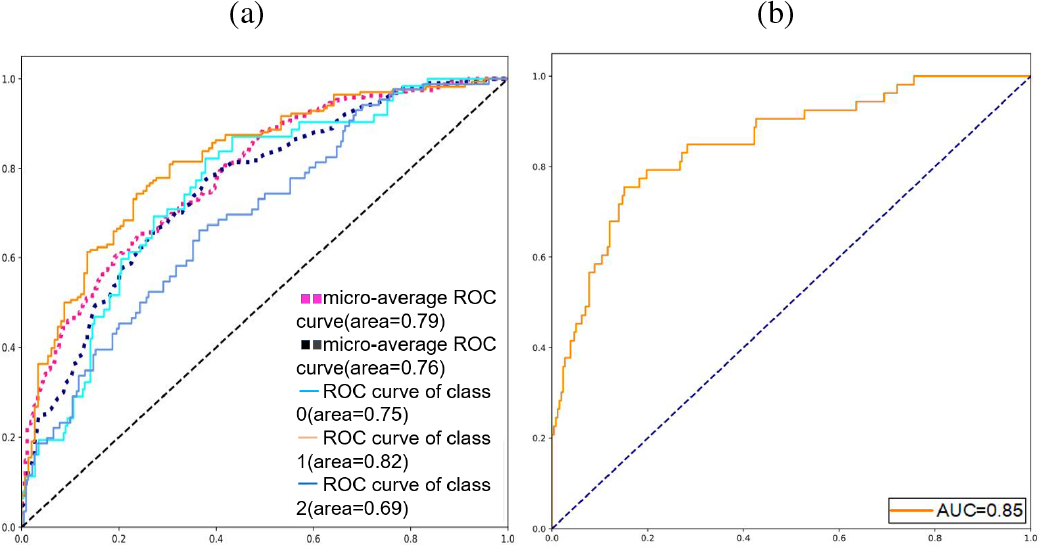
(a) ROC curves for three immune subtypes prediction by our method. (b) ROC curve achieved by using only pathological images to predict the IPS score representing the potential benefit from immune checkpoint inhibitor treatment.

Furthermore, to assess whether patients could derive benefit from immune checkpoint inhibitor treatment, we obtained the immunophenoscore (IPS) [44] from The Cancer Immunome Atlas that has analyzed the immune landscapes and antigenomes of 20 solid tumors. The IPS is calculated based on the gene expression of representative immune cells. A higher IPS score indicates a stronger immunogenicity and patient response to ICI treatment. As shown in Figure 6 (b), the average IPS score for the hot tumor subgroup was higher than that for the cold tumor subgroup, which also demonstrates the reliability of our method for classifying hot and cold tumors. So, we assigned each patient the label indicating the immunotherapy benefit based on the IPS scores. If the PD-1 IPS score increased compared to the control IPS score, the label of immunotherapy benefit is 1 and 0 otherwise. Subsequently, we used the pathological features to predict the immunotherapy benefit. The ROC-AUC value reached 0.85, indicated that our model can accurately predict benefit for patient from immune checkpoint inhibitor treatment using only pathological images.

### F. Contrastive learning and attentive pooling improve performance

We also conducted a comparative analysis of three selfsupervised learning models, namely masked autoencoder (MAE), momentum contrastive learning(MoCo) and adversarial contrastive learning (AdCo), regarding their performance in extracting tile-level features. To ensure a fair comparison, we fine-tuned the parameters of the three algorithms and utilized the extracted features to predict the level of immune cell infiltration. As depicted in Figure 7 (a), the results indicated that the performance of AdCo performed significantly superior to MAE and MoCo. Specifically, the predictive performance of AdCo-extracted features yielded a correlation coefficient of 0.30, while MAE and MoCo reached only 0.17 and 0.15. We speculated that the transformer-based [45] MAE is more inclined to capture global associations such as in natural images. In contrast, adversarial contrastive learning has the advantage in extracting local fine-grained feature in high-resolution pathology image, thereby resulted in better performance.

**Fig. 7:**
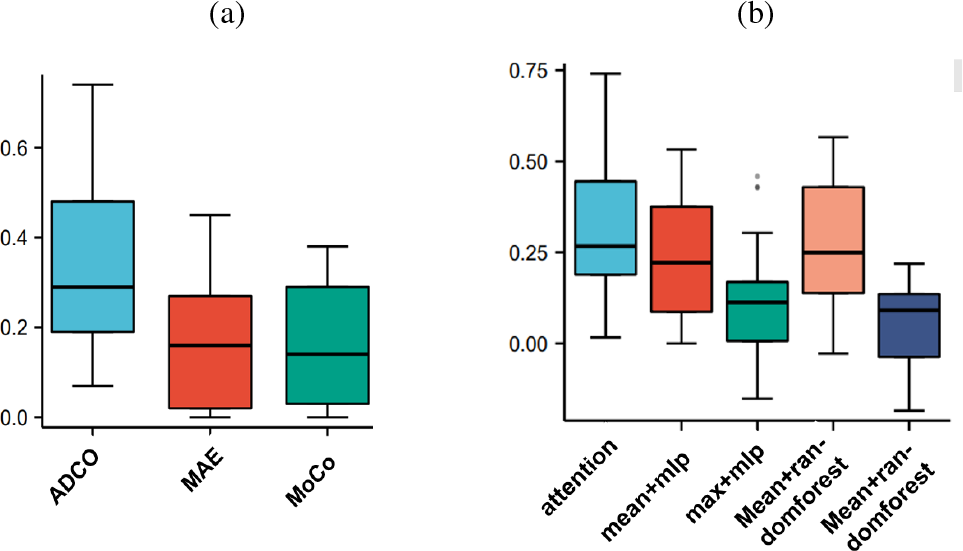
Model ablation experiments support that contrastive learning and attentive pooling improve performance. (a) Performance comparison in predicting tumor immune infiltration level by AdCo,MoCo and MAE feature extractors. (b) Comparison among different feature aggregation strategies and predictor for downstream task.

In addition, we examined the influence of tile-level feature aggregation strategies and predictors on the performance of downstream tasks. We compared several approaches, including attentive pooling + MLP, max pooling + MLP, average pooling + MLP, max pooling + RandomForest, and average pooling + RandomForest. All of these methods used tile-level features obtained from AdCo to predict immune cell infiltration levels. As depicted in Figure 7 (b), we discovered that the feature aggregation strategy had a significant impact on performance. Attention pooling yielded the best results, followed by average pooling, with max pooling performing the worst. The downstream predictor had a less significant effect in comparison.

### G. Spatial localization of tumor regions and immune cells via attention scores

In downstream tasks, we employed attentive pooling to aggregate tile-level features and acquired attention weights for each tile by weakly supervised learning, thereby endowing our model with a degree of interpretability. Initially, we evaluated the model capacity for spatial localization of tumor regions. For this purpose, we trained a tumor diagnosis model utilizing slide-level labels (tumor or normal) and cross-entropy as the loss function. As depicted in Figure 8 (a), the accuracy of tumor diagnosis reached 0.99 accuracy.

**Fig. 8:**
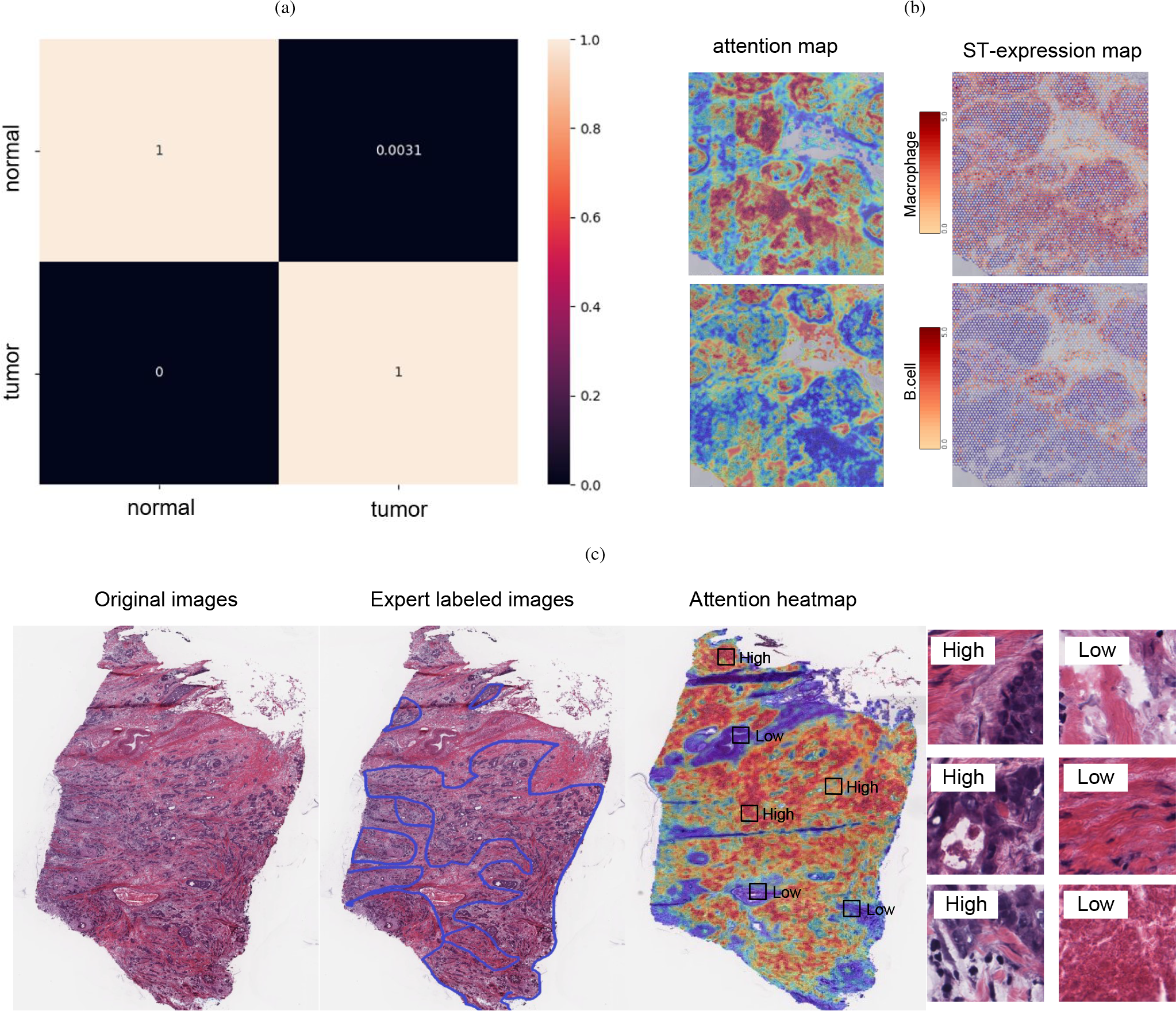
Tumor diagnosis and spatial localization of tumor regions via attention scores. (a) Confusion matrix for the tumor diagnosis task achieved by our model on TCGA-BRCA cohort. (b) Spatial expression landscapes of immune cell marker genes constructed by via attention scores and spatial transcriptomic data. Left column showed the heatmaps of the macrophages and B cells generated through the learned attention scores, and the right column showed the spatial expression landscape of the marker genes of these two cell types generated by spatial transcriptomic data. (c) Comparison of the spatial deconvolution of the tumor area indicated by attention scores with pathologist annotations. The heatmaps are generated by coloring each tile according to its attention score, with some representative tiles with highand low-attention scores.

Spatial transcriptomics measures the abundance of RNA in each spot at high spatial resolution, linking histological morphology with spatially located gene expression. This allows us to use spatial transcriptomics data to verify the ability of our model to locate specific immune cells spatially. We generated a heatmap using the tile attention scores generated from predicting macrophage infiltration levels and compared it to the macrophages spatial map generated using spatial transcriptomics data. The attention heatmap was highly consistent with the spatial transcriptomics map, as shown in Figure 8 (b). We also generated heatmaps for other immune cells and obtained similar results.

Finally, we generated an attention heatmap using coloring tiles using their attention scores and applying spatial deconvolution to the original pathology image. As shown in Figure 8 (c) and Figure S2(b), the areas with high-attention scores were highly consistent with the boundaries of tumor regions annotated by an experienced pathologist. Moreover, we visualized several tiles and noted that tiles with low attention scores predominantly comprised normal tissue while tiles with high attention scores largely encompassed tumor regions.

## IV. Discussion and Conclusion

In recent years, immunotherapy has received widespread attention due to its significant efficacy in the treatment of solid tumors. The tumor immune microenvironment plays an important role in the response to immunotherapy, and the development of effective biomarker based on TIME characteristics has become a focal point in the anti-tumor research community. High infiltration of cytotoxic lymphocytes are the main reason for the resistance of immunotherapy. Therefore, it is crucial to stratify patients who may benefit from immunotherapy and develop combinatorial treatment such as chemotherapy/radiotherapy and immunotherapy. However, due to technical and cost constraints, RNA sequencing-based tumor immune microenvironment has not been widely applied in clinical practice, especially in resource-limited settings. The advancement of computational pathology provides a new way to address this issue. We have demonstrated that the infiltration of immune cells into tumor can be effectively inferred from histological and morphological phenotype. The spatial deconvolution of pathological images greatly enhances thevisualization of the spatial localization of tumor and normal cells, providing assistance for personalized immunotherapy.

The superior performance of the model can be primarily attributed to the informative features extracted vis selfsupervised contrastive learning. This approach has proven to be particularly advantageous in the analysis of pathological images, where a vast quantity of unlabeled tiles can be directly utilized without the need for manual annotation. Notably, contrastive learning captures tile-level features more effectively, and the potential for self-supervised models in the field of pathological imaging warrants further exploration.

In summary, this paper proposed a deep learning method to infer the level of tumor immune cell infiltration from pathological images. Our experimental results demonstrate that pre-training based on contrastive learning effectively enhances the performance of downstream tasks. We believe that our study has made a significant contribution to advancing the field of computational pathology.

## Author contributions

H.L. and G.Y. conceived the study and designed the experiments. Y.Z. provided useful suggestions. G.Y. performed the experimental analysis. H.L. and G.Y. prepared the manuscript.

B.W. and H.L. revised the manuscript. B.W. and H.L. supervised the research.

## Acknowledgment

We are very grateful to Dr. Meihua Wang from the Department of Pathology at Suzhou University Affiliated Changzhou Cancer Hospital for his hard work to annotate the pathology images, as well as his valuable suggestion to our study.

## Code availability

All code was implemented using PyTorch as the primary deep-learning library. The complete pipeline for processing WSIs as well as training and evaluating the deep-learning models is available at https://github.com/hliulab/wsi2time.

## Guobang Yu

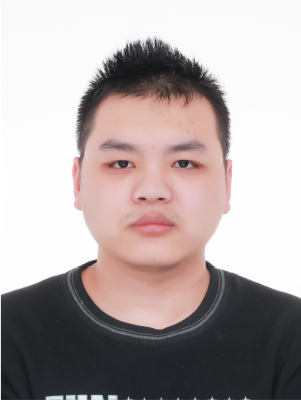

received his bachelor’s degree from Nanjing Forestry University. He is now a master student at College of Computer and Information Engineering, Nanjing Tech University, Nanjing, China. His interests include computational pathology and Bioinformatics.

## Yi Zuo

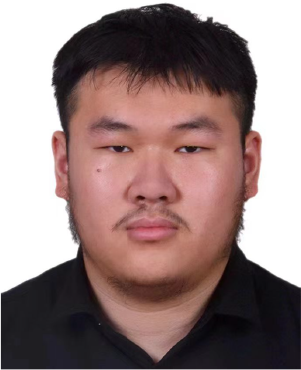

is an undergraduate student at College of Computer and Information Engineering, Nanjing Tech University, Nanjing, China. He is interested in medical image processing.

## Bin Wang

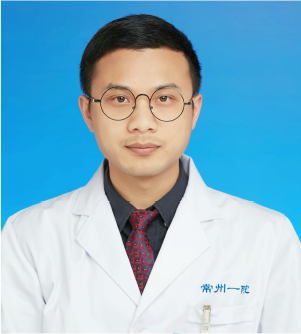

received his Ph.D. degree in cardiothoracic surgery from Fudan University, Shanghai, China, in 2016. He is currently a chief doctors at Department of Cardiothoracic Surgery, The third affiliated hospital of Soochow University, 213100, Changzhou, China. His research interests include Bioinformatics and Medical Imaging.

## Hui Liu

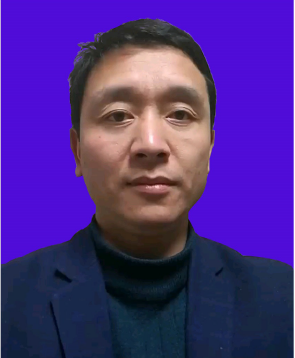

received his Ph.D. degree in computer science from Fudan University, Shanghai, China, in 2010. He is currently a professor at College of Computer and Information Engineering, Nanjing Tech University, Nanjing, China. His research interests include Medical Image, AI-driven drug discovery and Deep Learning. He has published dozens of papers in top-ranking conferences and journals.

## References

[1] J. Ferlay, M. Ervik, F. Lam, C. M, M. L, P. M, Z. A, S. I, and B. F, Global Cancer Observatory: Cancer Today, vol. 68. 2020.

[2] Y. Shiravand, F. Khodadadi, S. M. A. Kashani, S. R. Hosseini-Fard, S. Hosseini, H. Sadeghirad, R. Ladwa, K. O’byrne, and A. Kulasinghe, “Immune checkpoint inhibitors in cancer therapy,” Current Oncology, vol. 29, pp. 3044–3060, 2022.

[3] Z. Wang, K. Sun, Y. Xiao, B. Feng, K. Mikule, X. Y. Ma, N. Feng, C. P. Vellano, L. Federico, J. R. Marszalek, G. B. Mills, J. Hanke, S. Ramaswamy, and J. Wang, “Niraparib activates interferon signaling and potentiates anti-pd-1 antibody efficacy in tumor models,” Scientific Reports, vol. 9, p. 1853, 2019.

[4] Q. Xu, S. Chen, Y. Hu, and W. Huang, “Landscape of immune microenvironment under immune cell infiltration pattern in breast cancer,” Frontiers in Immunology, vol. 12, p. 711433, 2021.

[5] F. Tang, C. Barbacioru, W. Yangzhou, E. Nordman, C. Lee, N. Xu, X. Wang, J. Bodeau, B. B. Tuch, A. Siddiqui, K. Lao, M. A. Surani, et al., “mrna-seq whole-transcriptome analysis of a single cell,” Nature Methods, vol. 6, p. 377–382, 2009.

[6] A. M. Newman, C. B. Steen, C. L. Liu, A. J. Gentles, A. A. Chaudhuri, F. Scherer, M. S. Khodadoust, M. S. Esfahani, B. A. Luca, D. Steiner, M. Diehn, and A. A. Alizadeh, “Robust enumeration of cell subsets from tissue expression profiles,” Nat Methods, vol. 12, pp. 453–457, 2015.

[7] A. Subramanian, P. Tamayo, V. K. Mootha, S. Mukherjee, B. L. Ebert, M. A. Gillette, A. Paulovich, S. L. Pomeroy, T. R. Golub, E. S. Lander, and J. P. Mesirov, “Gene set enrichment analysis: A knowledge-based approach for interpreting genome-wide expression profiles,” Proceedings of the National Academy of Sciences, vol. 102, no. 43, pp. 15545–15550, 2005.

[8] K. Yoshihara, M. Shahmoradgoli, E. Martínez, et al., “Inferring tumour purity and stromal and immune cell admixture from expression data,” Nature Communications, vol. 4, p. 2612, 2013.

[9] L. Yann, B. Yoshua., and H. Geoffrey, “Deep learning,” Nature, vol. 521, p. 436–444, 2015.

[10] J. Deng, W. Dong, R. Socher, L.-J. Li, K. Li, and L. Fei-Fei, “Imagenet: A large-scale hierarchical image database,” in 2009 IEEE Conference on Computer Vision and Pattern Recognition, pp. 248–255, 2009.

[11] A. Krizhevsky, I. Sutskever, and G. E. Hinton, “Imagenet classification with deep convolutional neural networks,” Commun. ACM, vol. 60, p. 84–90, may 2017.

[12] K. Simonyan and A. Zisserman, “Very deep convolutional networks for large-scale image recognition,” CoRR, vol. abs/1409.1556, 2014.

[13] K. He, X. Zhang, S. Ren, and J. Sun, “Deep residual learning for image recognition,” in 2016 IEEE Conference on Computer Vision and Pattern Recognition (CVPR), pp. 770–778, 2016.

[14] S. Wang, R. Rong, D. M. Yang, J. Fujimoto, S. Yan, L. Cai, L. Yang, D. Luo, C. Behrens, E. R. Parra, B. Yao, L. Xu, T. Wang, X. Zhan, I. I. Wistuba, J. Minna, Y. Xie, and G. Xiao, “Computational staining of pathology images to study the tumor microenvironment in lung cancer,” Cancer Research, vol. 80, p. 2056–2066, 2020.

[15] G. Campanella, M. G. Hanna, L. Geneslaw, A. Miraflor, V. W. K. Silva, K. J. Busam, E. Brogi, V. E. Reuter, D. S. Klimstra, and T. J. Fuchs, “Clinical-grade computational pathology using weakly supervised deep learning on whole slide images,” Nature Medicine, vol. 25, p. 1301–1309, 2019.

[16] M. Ilse, J. M. Tomczak, and M. Welling, “Chapter 22 - deep multiple instance learning for digital histopathology,” in Handbook of Medical Image Computing and Computer Assisted Intervention (S. K. Zhou, D. Rueckert, and G. Fichtinger, eds.), The Elsevier and MICCAI Society Book Series, pp. 521–546, Academic Press, 2020.

[17] A. M. Newman, C. L. Liu, M. R. Green, A. J. Gentles, W. Feng, Y. Xu, C. D. Hoang, M. Diehn, and A. A. Alizadeh, “Robust enumeration of cell subsets from tissue expression profiles,” Nature Methods, vol. 12, pp. 453–457, 2015.

[18] C. L. Chen, C. C. Chen, W. H. Yu, S. H. Chen, Y. C. Chang, T. I. Hsu, M. Hsiao, C. Y. Yeh, and C. Y. Chen, “An annotation-free wholeslide training approach to pathological classification of lung cancer types using deep learning,” Nature Communications, vol. 12, p. 1193, 2021.

[19] M. Y. Lu, D. F. Williamson, T. Y. Chen, R. J. Chen, M. Barbieri, and F. Mahmood, “Data-efficient and weakly supervised computational pathology on whole-slide images,” Nature Biomedical Engineering, vol. 5, p. 555–570, 2021.

[20] J. N. Kather, A. T. Pearson, N. Halama, D. Jäger, J. Krause, S. H. Loosen, A. Marx, P. Boor, F. Tacke, U. P. Neumann, H. I. Grabsch, T. Yoshikawa, H. Brenner, J. Chang-Claude, M. Hoffmeister, C. Trautwein, and T. Luedde, “Deep learning can predict microsatellite instability directly from histology in gastrointestinal cancer,” Nature Medicine, vol. 25, p. 1054–1056, 2019.

[21] J. N. Kather, L. R. Heij, H. I. Grabsch, C. Loeffler, A. Echle, H. S. Muti, J. Krause, J. M. Niehues, K. A. Sommer, P. Bankhead, L. F. Kooreman, J. J. Schulte, N. A. Cipriani, R. D. Buelow, P. Boor, N. Ortiz-Brüchle, A. M. Hanby, V. Speirs, S. Kochanny, A. Patnaik, A. Srisuwananukorn, H. Brenner, M. Hoffmeister, P. A. van den Brandt, D. Jäger, C. Trautwein, A. T. Pearson, and T. Luedde, “Pan-cancer image-based detection of clinically actionable genetic alterations,” Nature Cancer, vol. 1, p. 789–799, 2020.

[22] J. Hu, X. Li, K. Coleman, A. Schroeder, N. Ma, D. J. Irwin, E. B. Lee, R. T. Shinohara, and M. Li, “Spagcn: Integrating gene expression, spatial location and histology to identify spatial domains and spatially variable genes by graph convolutional network,” Nature Methods, vol. 18, p. 1342–1351, 2021.

[23] B. Schmauch, A. Romagnoni, E. Pronier, C. Saillard, P. Maillé, J. Calderaro, A. Kamoun, M. Sefta, S. Toldo, M. Zaslavskiy, T. Clozel, M. Moarii, P. Courtiol, and G. Wainrib, “A deep learning model to predict rna-seq expression of tumours from whole slide images,” Nature Communications, vol. 11, p. 1193, 2020.

[24] S. Y. Yoo, H. E. Park, J. H. Kim, X. Wen, S. Jeong, N. Y. Cho, H. G. Gwon, K. Kim, H. S. Lee, S. Y. Jeong, K. J. Park, S. W. Han, T. Y. Kim, J. M. Bae, and G. H. Kang, “Whole-slide image analysis reveals quantitative landscape of tumor-immune microenvironment in colorectal cancers,” Clinical Cancer Research, vol. 26, p. 870–881, 2020.

[25] J. Saltz, R. Gupta, L. Hou, T. Kurc, P. Singh, and et al., “Spatial organization and molecular correlation of tumor-infiltrating lymphocytes using deep learning on pathology images,” Cell Reports, vol. 23, pp. 181–193, 2018.

[26] A. v. d. Oord, Y. Li, and O. Vinyals, “Representation learning with contrastive predictive coding,” arXiv preprint arXiv:1807.03748, 2018.

[27] K. He, H. Fan, Y. Wu, S. Xie, and R. Girshick, “Momentum contrast for unsupervised visual representation learning,” in Proceedings of the IEEE/CVF conference on computer vision and pattern recognition, pp. 9729–9738, 2020.

[28] H. Huang, G. Zhou, X. Liu, L. Deng, C. Wu, D. Zhang, and H. Liu, “Contrastive learning-based computational histopathology predict differential expression of cancer driver genes,” Briefings in Bioinformatics, vol. 23, 07 2022. bbac294.

[29] Z.-H. Zhou, “A brief introduction to weakly supervised learning,” National Science Review, vol. 5, pp. 44–53, 08 2017.

[30] M. Ilse, J. Tomczak, and M. Welling, “Attention-based deep multiple instance learning,” in International conference on machine learning, pp. 2127–2136, PMLR, 2018.

[31] S. Hong, Y. Zou, and W. Wang, “Gated multi-head attention pooling for weakly labelled audio tagging,” Proceedings of the Annual Conference of the International Speech Communication Association, INTERSPEECH, vol. 2020-October, pp. 816–820, 2020.

[32] N. Otsu, “A threshold selection method from gray-level histograms,” IEEE Transactions on Systems, Man, and Cybernetics, vol. 9, no. 1, pp. 62–66, 1979.

[33] Q. Hu, X. Wang, W. Hu, and G.-J. Qi, “Adco: Adversarial contrast for efficient learning of unsupervised representations from self-trained negative adversaries,” in Proceedings of the IEEE/CVF Conference on Computer Vision and Pattern Recognition, pp. 1074–1083, 2021.

[34] K. He, X. Chen, S. Xie, Y. Li, P. Dollár, and R. Girshick, “Masked autoencoders are scalable vision learners,” in Proceedings of the IEEE/CVF conference on computer vision and pattern recognition, pp. 16000– 16009, 2022.

[35] M. Tekguc, J. B. Wing, M. Osaki, J. Long, and S. Sakaguchi, “Treg-expressed ctla-4 depletes cd80/cd86 by trogocytosis, releasing free pd-l1 on antigen-presenting cells,” Proceedings of the National Academy of Sciences, vol. 118, no. 30, p. e2023739118, 2021.

[36] X. Cao, B. Li, J. Chen, J. Dang, S. Chen, E. G. Gunes, B. Xu, L. Tian, S. Muend, M. Raoof, C. Querfeld, J. Yu, S. T. Rosen, Y. Wang, and M. Feng, “Effect of cabazitaxel on macrophages improves cd47targeted immunotherapy for triple-negative breast cancer,” Journal for ImmunoTherapy of Cancer, vol. 9, no. 3, 2021.

[37] C. Yunna, H. Mengru, W. Lei, and C. Weidong, “Macrophage m1/m2 polarization,” European Journal of Pharmacology, vol. 877, p. 173090, 2020.

[38] Q. Xu, S. Chen, Y. Hu, and W. Huang, “Clinical m2 macrophages-related genes to aid therapy in pancreatic ductal adenocarcinoma,” Cancer Cell International, vol. 21, p. 582, 2021.

[39] Z. Q. Wang, K. Milne, J. R. Webb, and P. H. Watson, “Cd74 and intratumoral immune response in breast cancer,” Oncotarget, vol. 8, pp. 12664–12674, 2017.

[40] L. Liao, W. Dang, T. Lin, J. Yu, T. Liu, W. Li, S. Xiao, L. Feng, J. Huang, R. Fu, J. Li, L. Liu, M. Wang, H. Tao, H. Jiang, K. Chen, X. Diao, B. Zhou, X. Shen, and C. Luo, “A potent pgk1 antagonist reveals pgk1 regulates the production of il-1 and il-6,” Acta Pharmaceutica Sinica B, vol. 12, no. 11, pp. 4180–4192, 2022.

[41] N. J. Edwards, M. Oberti, R. R. Thangudu, S. Cai, P. B. McGarvey, S. Jacob, S. Madhavan, and K. A. Ketchum, “The cptac data portal: A resource for cancer proteomics research,” Journal of Proteome Research, vol. 14, p. 2707–2713, 2015.

[42] A. Mlynska, R. Vaišnorė, V. Rafanavičius, S. Jocys, J. Janeiko, M. Petrauskytė, S. Bijeikis, P. Cimmperman, B. Intaitė, K. Žilionytė, A. Barakauskienė, R. Meškauskas, E. Paberalė, and V. Pašukonienė, “A gene signature for immune subtyping of desert, excluded, and inflamed ovarian tumors,” American Journal of Reproductive Immunology, vol. 84, p. e13244, 2020.

[43] J. Galon and D. Bruni, “Approaches to treat immune hot, altered and cold tumours with combination immunotherapies,” Nature Reviews Drug Discovery, vol. 18, p. 197–218, 2019.

[44] P. Charoentong, F. Finotello, M. Angelova, C. Mayer, M. Efremova, D. Rieder, H. Hackl, and Z. Trajanoski, “Pan-cancer immunogenomic analyses reveal genotype-immunophenotype relationships and predictors of response to checkpoint blockade,” Cell Reports, vol. 18, pp. 248–262, 2017.

[45] A. Vaswani, N. Shazeer, N. Parmar, J. Uszkoreit, L. Jones, A. N. Gomez, Ł. Kaiser, and I. Polosukhin, “Attention is all you need,” Advances in neural information processing systems, vol. 30, 2017.

